# Factorbook: an Updated Catalog of Transcription Factor Motifs and Candidate Regulatory Motif Sites

**DOI:** 10.1101/2021.10.11.463518

**Authors:** Henry E. Pratt, Gregory R. Andrews, Nishigandha Phalke, Michael J. Purcaro, Arjan van der Velde, Jill E. Moore, Zhiping Weng

## Abstract

The human genome contains roughly 1,600 transcription factors (TFs) (1), DNA-binding proteins recognizing characteristic sequence motifs to exert regulatory effects on gene expression. The binding specificities of these factors have been profiled both *in vitro*, using techniques such as HT-SELEX (2), and *in vivo*, using techniques including ChIP-seq (3, 4). We previously developed Factorbook, a TF-centric database of annotations, motifs, and integrative analyses based on ChIP-seq data from Phase II of the ENCODE Project. Here we present an update to Factorbook which significantly expands the breadth of cell type and TF coverage. The update includes an expanded motif catalog derived from thousands of ENCODE Phase II and III ChIP-seq experiments and HT-SELEX experiments; this motif catalog is integrated with the ENCODE registry of candidate cis-regulatory elements to annotate a comprehensive collection of genome-wide candidate TF binding sites. The database also offers novel tools for applying the motif models within machine learning frameworks and using these models for integrative analysis, including annotation of variants and disease and trait heritability. We will continue to expand the resource as ENCODE Phase IV data are released.

## INTRODUCTION

The human genome includes the instructions for producing an estimated 1,600 transcription factors (TFs), a broad class of proteins that interact with DNA in order to modulate regulatory element activity and gene expression (1). TFs typically possess a DNA binding domain (DBD) which recognizes 6-20 base-pair (bp) long characteristic consensus binding sequences, or a *motif*, present within the TF’s target regulatory elements. TFs may be grouped according to several known families of DBDs which frequently recognize similar DNA sequences.

Numerous resources have been developed to catalog TF motifs. The HOCOMOCO catalog (5) indexes binding specificities for nearly 700 human TFs and more than 400 mouse TFs identified from ChIP-seq and HT-SELEX data, and the JASPAR catalog (6) contains more than 700 curated non-redundant binding profiles for eukaryotic TFs. The earlier UniPROBE (7) resource contains more than 700 binding profiles from *in vitro* protein binding microarray experiments, and the broader CisBP (8) incorporates data from these sources and others, including our previous release of Factorbook (9, 10), to annotate both measured and inferred binding profiles for thousands of TFs across tens of species.

Here we present an update to Factorbook which leverages the extensive ChIP-seq data available through Phase III of the ENCODE Project to build a comprehensive TF motif catalog for more than 1,000 TFs. We also provide two notable features not available in existing catalogs to our knowledge. First, we catalog motif models built using convolutional neural networks (CNNs), which are finding increasing applications in genomics including in discovering TF motifs and predicting TF binding (11–13); these will be easily integrated into future models for transfer learning. Second, we leverage the ENCODE Registry of candidate Cis-Regulatory Elements (14) to provide a genome-wide catalog of motif sites in regulatory sequences, with associated epigenetic and evolutionary annotations; we illustrate the usefulness of this catalog for downstream applications by using it to quantify trait heritability using partitioned LD score regression (15).

## OVERVIEW

Factorbook is a transcription factor-centric database cataloging information for 694 distinct human TFs and 62 mouse TFs profiled in 249 and 38 human and mouse cell types; this is a substantial increase from the 119 human TFs in the first Factorbook release (10). At this scale, Factorbook’s previous matrix-view entry point (10) has become intractable; the primary entry point is now a factor search (**Figure 1A**), which directs users to a detailed information page for the searched TF. Each factor’s information page contains curated information from various external sources, including NCBI, Uniprot, HGNC, and Ensembl (**Figure 1B**). Rich data tables separately listing the available TFs and cell types are also available for browsing (**Figure 1C**). Furthemore, we display the expression levels of each TF using ENCODE RNA-seq data in a variety of primary cells, primary tissues, and cell lines (**Figure 1D**), and also display primary data resulting from several integrative analyses described in detail below.

**Figure 1.**
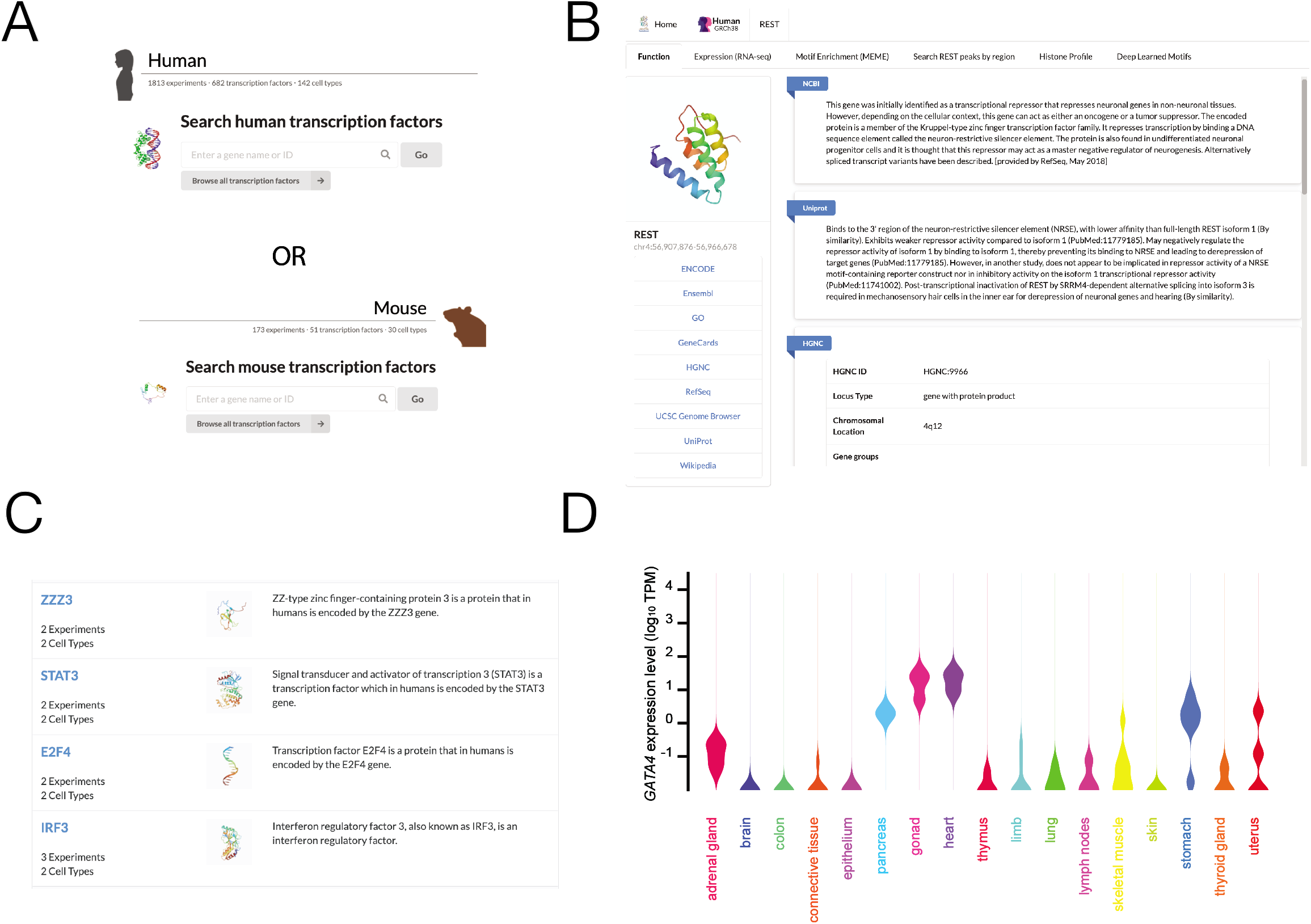
Overview of the main Factorbook interface. (**A**) An example of the main TF search for human and mouse. (**B**) The information page for REST, highlighting information curated from external sources. (**C**) The transcription factor table, listing all 682 human TFs with available data for browsing. (**D**) Factorbook’s display of the RNA-seq expression profile for *GATA4* in human embryonic primary tissues.

From a technical perspective, we have replaced the Wiki-based technology of the first Factorbook release (10) with an architecture utilizing a ReactJS frontend and GraphQL Application Programming Interfaces (APIs). This offers a number of improvements and novel capabilities, including 1) facilitating programmatic data access, 2) offering interactive rather than static visualizations, and 3) enabling Factorbook to perform a variety of interactive analyses described in detail below, including SNP annotation and intersection of resources with user-uploaded BED files in real time.

### A comprehensive motif catalog derived from ChIP-seq and HT-SELEX data

A cornerstone of the primary data contained within Factorbook is a comprehensive catalog of TF recognition motifs. We expanded on previous catalogs (5, 16) in two ways: first, we aimed to annotate binding sequences for as many transcription factors as possible, including TFs not profiled by other efforts to our knowledge (5, 8, 17, 18); and second, we aimed to provide motifs in optimal formats that will integrate seamlessly into the variety of machine learning frameworks which are actively being developed to study TF binding (12, 13, 19) in addition to conventional downstream analysis using tools such as the MEME suite (20).

We thus designed two complementary pipelines for de novo motif identification (**Methods**). First, we applied our previous MEME-based pipeline (9) to the top 500 strongest ChIP-seq peaks from each transcription factor ChIP-seq dataset produced during the first three phases of ENCODE. This pipeline identifies up to five enriched motifs per ChIP-seq dataset; these motifs are subsequently filtered by quality control metrics we developed previously (9), including peak centrality and enrichment against permuted genomic sequences. Second, we developed a convolutional neural network, *ZMotif*, for motif discovery applicable to HT-SELEX (**Methods**).

In total, MEME identifies 6,921 motifs from human ChIP-seq datasets. The MEME catalog contains several redundant motifs for well-profiled factors such as CTCF and REST; we therefore applied UMAP (21, 22) to the motifs to map them into a reduced-dimension space. The UMAP projection for MEME motifs is shown in **Figure 2A**, with several clusters of known motifs annotated. The number of motif clusters ranges from 100 to 300 depending on hyperparameter selection; we make several interactive UMAP plots with different hyperparameters available through Factorbook to aid users in identifying motif clusters for downstream analysis. We then applied TOMTOM (23) to compare our MEME motifs against the HOCOMOCO and JASPAR catalogs (5, 16); we find that the Factorbook catalog includes nearly 100% of the motifs in these two sources, and further identifies novel motifs not present therein; novel motifs include candidate motifs for 358 TFs which are not profiled in HOCOMOCO or JASPAR to our knowledge, including 101 of 428 factors previously classified as “likely sequence-specific factors” (1) but without annotated binding specificity annotated elsewhere (**Figure 2B**). One example novel motif is that of ZNF407 (**Figure 2C**), which shows high evolutionary conservation (**Figure 2D**), prefers to reside in the center of ChIP-seq peaks (**Figure 2E**), and is protected from DNase I cleavage (**Figure 2F**). Each individual factor page contains indexed lists of all motifs identified by MEME; these and the HT-SELEX motifs can also be searched through Factorbook either by consensus sequence or by uploading motifs in MEME format to match against the catalog; visual results are provided in real time (**Figure 2G**).

**Figure 2.**
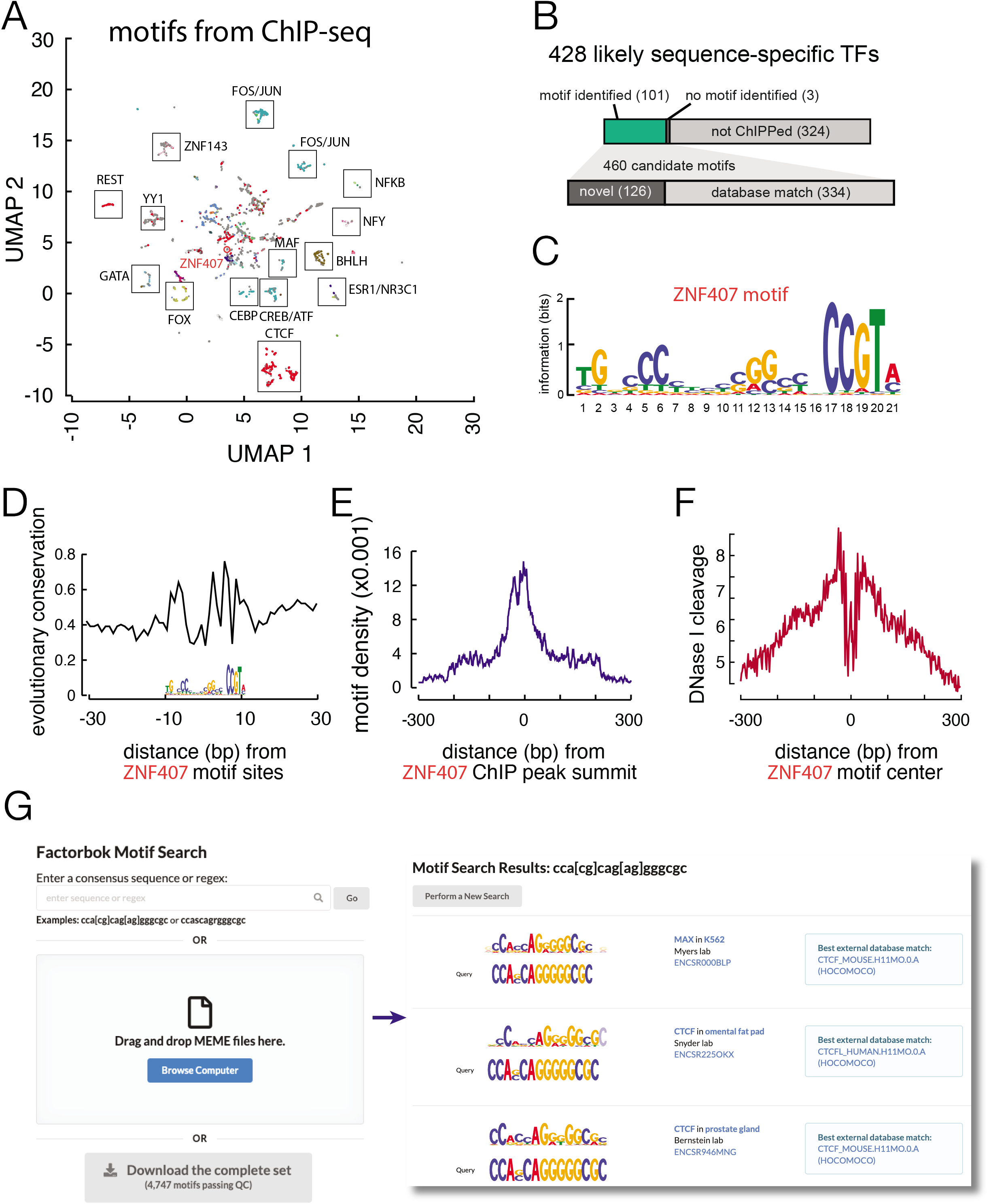
Overview of the Factorbook MEME ChIP-seq motif catalog. (**A**) A UMAP projection of motifs passing QC, with some DNA binding domains colored (C2H2 Zinc Finger in red, GATA in teal, nuclear receptor in dark purple) and several motif clusters annotated. (**B**) Overview of novel motifs cataloged by Factorbook. (**C**) A novel motif for ZNF407, with supporting evolutionary conservation (phyloP 100-way) (**D**), peak centrality (**E**), and DNase-seq footprint (**F**) aggregate plots. (**G**) The Factorbook motif search interface, showing matches for the consensus sequence for the CTCF motif.

ZMotif identifies a total of 6,700 motifs from HT-SELEX two public HT-SELEX datasets (2, 24); we perform similar UMAP projection and cluster annotation for these motifs (**Figure 3A**). SELEX motifs are also available for visualization on each TF’s information page; motifs are shown for each HT-SELEX cycle, along with enrichment statistics (**Figure 3B**). Motifs from all three methods are made available for download as PWMs in MEME format; deep-learned filters are further available in HDF5 format that can be directly used by models such as neural networks for transfer learning. The motif page also displays TOMTOM matches against HOCOMOCO and JASPAR for each motif.

**Figure 3.**
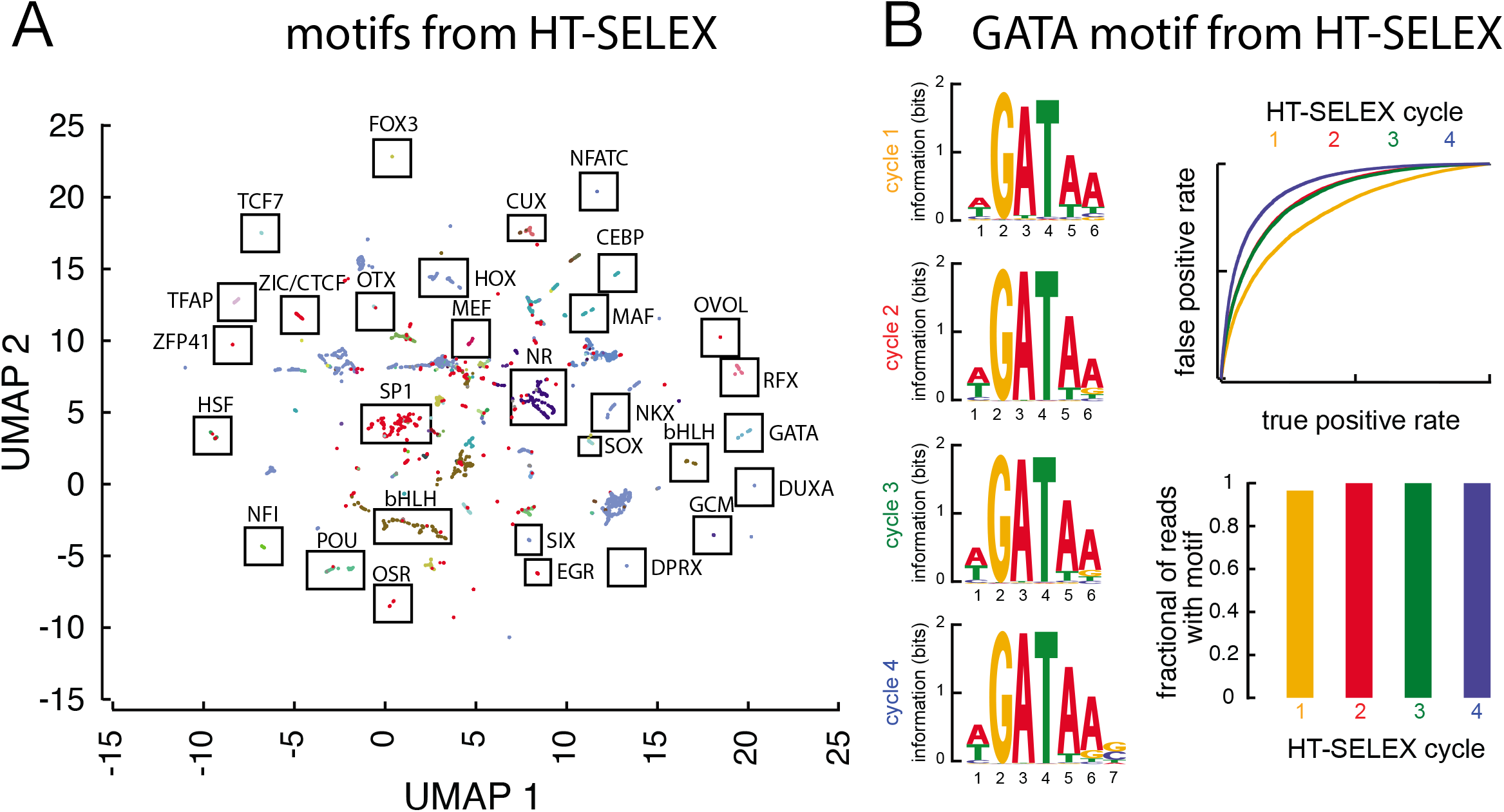
The Factorbook HT-SELEX motif catalog. (**A**) A UMAP projection of 6,700 HT-SELEX motifs with clusters annotated. (**B**) Example of the SELEX motif interface for GATA4, showing motifs for each of the four SELEX cycles for a GATA4 HT-SELEX experiment as well as two motif enrichment metrics, an ROC curve (top) and a readout of the fraction of reads at each cycle containing the motif (bottom).

### Genome-wide instances of motifs in ChIP-seq peaks

For ChIP-seq motifs identified with MEME, we use the FIMO tool from the MEME suite (20) to scan IDR thresholded ChIP-seq peaks from human TF ChIP-seq datasets for motif instances, filtering at the standard p-value cutoff of 10^−4^. We identified 110,001,176 (overlapping allowed) motif instances in total; when overlapping regions are merged, the motif sites number 6,720,871. These instances are available for download through Factorbook in BED format through the associated motif page; additionally, we have implemented a novel database-backed service allowing users to upload their own BED files for real-time intersection with motif instances, accessible through the same page. Bulk downloads of the complete set of BED files are also offered in TAR format.

To aid the user in evaluating how likely a given motif we identify is to be recognized by the TF of a ChIP-seq experiment (as opposed to, for example, a cofactor), we compute three metrics for the motif instances. First, we assess the evolutionary conservation of the motif instance and surrounding peak sequence using several phyloP scores across collections of vertebrates and mammals (25) (**Figure 2D**). Second, we compute the distance between each motif instance and the corresponding peak summit since the instances of bona fide motifs tend to be near peak summits (**Figure 2E**). Third, we compute the distribution of DNase-seq and ATAC-seq reads around each motif instance when matched data are available in the corresponding cell type (**Figure 2F**). We display histograms and aggregated signal profiles for each of these three metrics on the motif page. In general, we find that high quality motif instances are central within peaks, are more evolutionarily conserved than the surrounding peak sequences, and are less accessible to DNase I and Tn5 than surrounding non-motif peak sequences. Conservation scores are also available for individual motif sites in the motif site BED files, and the visualizations of the aggregate plots can also be exported in image or raw data formats.

### Genome-wide motif instances in candidate *cis*-regulatory elements

We previously developed the ENCODE Registry of candidate cis-regulatory elements (cCREs), a collection of nearly 1 million candidate human enhancers, promoters, and insulators, which are the subset of representative DNase hypersensitive sites (rDHSs) with high signals from two histone modifications (H3K4me3, a promoter mark, and H3K27ac, an enhancer mark) and the insulator-binding protein CTCF (14). The Registry integrates data from more than 1,000 cell types, while the transcription factor ChIP-seq data included in Factorbook derives primarily from five human cell lines (HepG2, K562, HEK293, GM12878, and MCF-7). Accordingly, motif instances in ChIP-seq peaks are most common within cCREs and rDHSs from the Registry that are active in primary cell types biologically similar to the aforementioned five cell lines. For example, embryonic bone marrow and liver are responsible for hematopoiesis, and nearly twice as many rDHSs active in those embryonic tissues contain peak motif instances as those active in other tissues, in line with the prevalence of ChIP-seq data in the red blood cell precursor K562 (**Figure 4A**). Therefore, we applied FIMO to identify instances of all the high-quality motifs from both our ChIP-seq MEME catalog and our HT-SELEX ZMotif catalog within cCREs and rDHSs. Given the larger scale of the rDHS set, we used a more stringent FIMO p-value threshold of 10^−6^ for MEME ChIP-seq motifs and 10^−5^ for ZMotif HT-SELEX motifs to reduce false positives. We also generated sets at more stringent thresholds of p<10^−7^ and p<10^−8^ for users preferring even higher confidence sets (**Figure 4B**).

**Figure 4.**
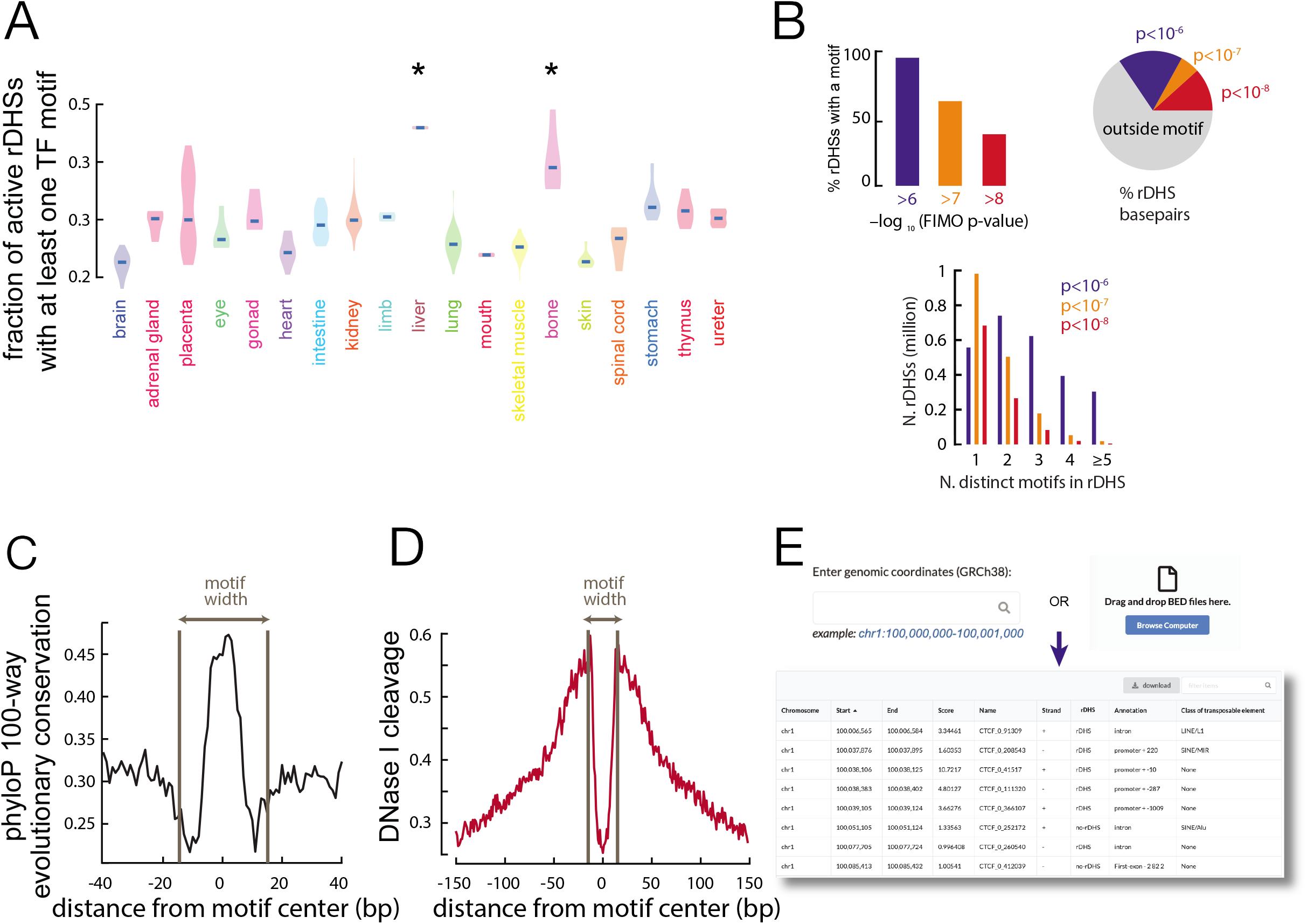
Overview of the Factorbook regulatory motif site catalog. (**A**) Fraction of representative DNase hypersensitive sites (rDHSs) active in a variety of embryonic cell types containing at least one motif identified in a ChIP-seq peak; cell types similar to K562 (fetal liver) and GM12878 (fetal bone) are most enriched due to the prevalence of ChIP-seq for those cell lines. (**B**) Distribution of motif sites within rDHSs and fraction of rDHS sequence covered by motifs at different thresholds. (**C**) Aggregated evolutionary conservation and (**D**) DNase I cleavage in embryonic kidney at 10,000 randomly chosen motif sites from the catalog within rDHSs active in embryonic kidney. (**E**) The motif site search interface, showing CTCF motif sites in a genomic region on chromosome 1.

In total, we define a catalog of 33,452,885 (overlapping allowed) candidate regulatory transcription factor motif sites within rDHSs using MEME-identified ChIP-seq motifs at a FIMO p-value of <10^−6^. These number 7,189,228 when overlapping motif sites are merged; of these, 4,902,200 (68.2%) are not present in the TF ChIP-seq peak motif site catalog. More than 95% of rDHSs contain at least one of these motif sites, with most having between 1 and 4; in total, the motif sites cover 30% of the base pairs in rDHS sequence (**Figure 4B**). The more stringent sets, numbering 2,840,049 and 1,572,634 non-overlapping motifs, respectively, cover a smaller portion of sequence within a smaller number of rDHSs (**Figure 4B**). We aggregate evolutionary conservation and DNase-seq reads at each of the motif sites at the most lenient threshold of p<10^−6^; this highlights that these motifs are significantly more conserved than surrounding rDHS sequences and are also less accessible to DNase I, suggesting the protection of the associated DNA by the bound TF. These findings regarding conservation (**Figure 4C**) and DNase I protection (**Figure 4D**) hold even for motifs present in rDHSs but not ChIP-seq peaks, supporting the idea that at least a subset of these motifs are true transcription factor binding sites which would be identified if ChIP-seq were performed in the correct biological context. These metrics are available through the motif page for each TF, as are the complete sets of instances in BED format. Instances can also be searched by BED file upload (**Figure 4E**). We performed the same analysis for HT-SELEX, identifying 9,205,043 distinct non-overlapping motif sites within rDHSs at a FIMO p-value <10-5, also accounting for roughly 30% of rDHS sequence. These sites are also available for download and searching through Factorbook.

### Tools for integrating motifs with GWAS results

It is hypothesized that many non-coding disease-associated variants confer risk for a given trait or disease by impacting transcription factor recognition sequences within regulatory elements. We therefore designed an interactive platform within Factorbook to facilitate the annotation of SNPs with candidate impacts on TF motif instances in our catalog, available through the Factorbook homepage. Users input a SNP’s rsID and optionally select a population or subpopulation from the 1,000 Genomes Project from which to include SNPs in linkage disequilibrium (LD) (**Figure 5A**). Factorbook intersects these SNPs with motif instances from our catalog in real time and displays the results, sorted by impact on the position weight matrix (PWM) of the canonical motif from MEME and the predicted impact on binding from our deep learning models (**Figure 5B**). Simultaneously, Factorbook searches all annotated TF peaks from ENCODE that intersect the SNPs, allowing users to determine if there is direct ChIP-seq support for any candidate TFs identified by motif analysis (**Figure 5C**).

**Figure 5.**
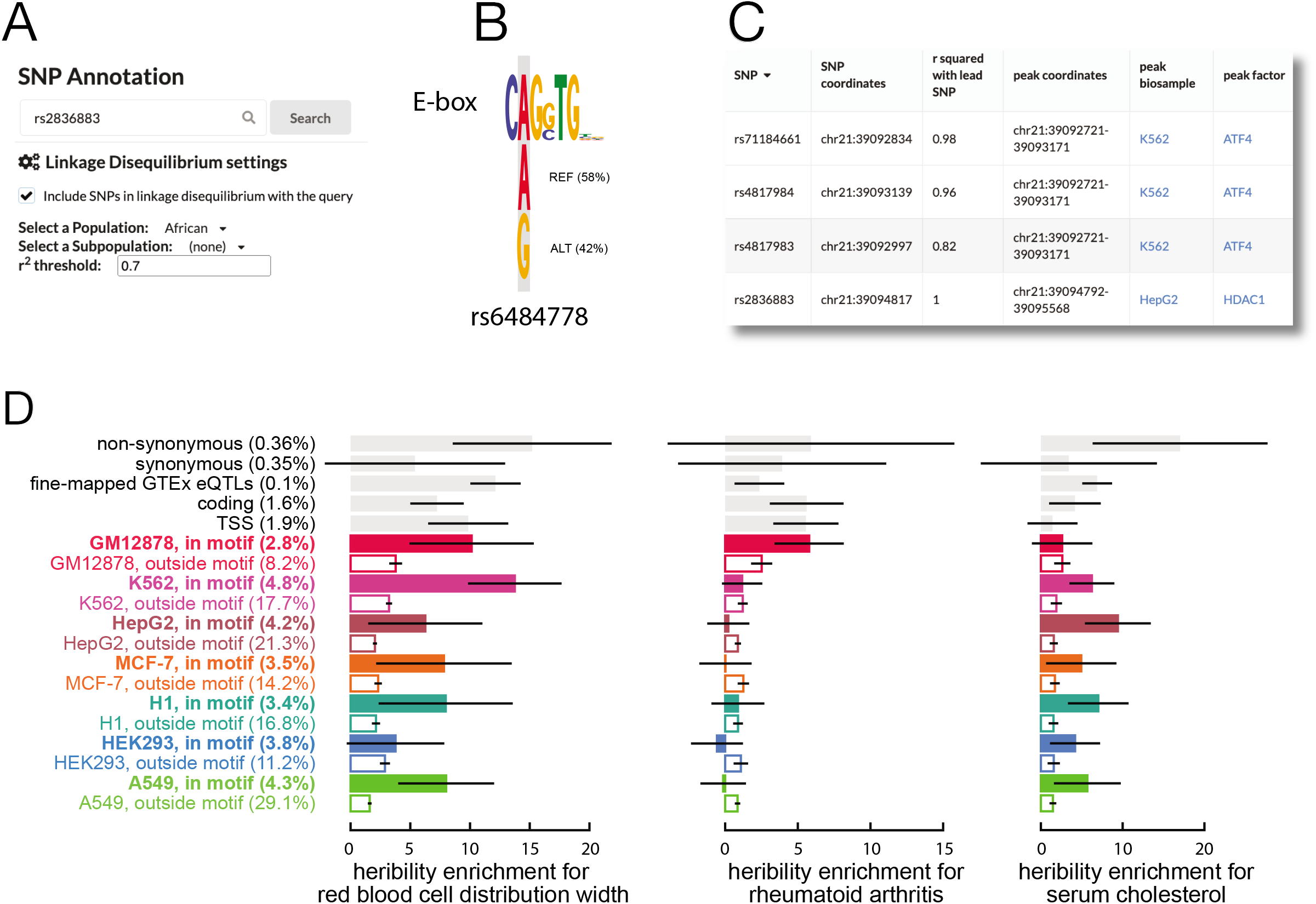
Annotation of variants with Factorbook. (**A**) The variant annotation interface; the user inputs a SNP rsID and can optionally select to include SNPs in LD with the searched SNP. (**B**) Example search results for a given SNP showing an impact on an E-box motif from the Factorbook rDHS motif site catalog. (**C**) The variant peak intersection view showing ChIP-seq peaks intersecting variants in LD with an example query. (**D**) Heritability enrichment for a variety of traits within motif sites identified in ChIP-seq peaks from seven distinct ENCODE cell lines, computed using partitioned LD score regression.

Additionally, we have built heritability models for partitioned LD score regression (15) from the motif sites in our catalog within ChIP-seq peaks. We provide one model which includes the complete set of motif instances as well as one which includes motif sites grouped by the cell type in which the corresponding ChIP-seq peak was identified. Overall, heritability for traits is highly enriched within motif sequences, with enrichment generally being the strongest within motif sites identified in ChIP-seq peaks from disease-relevant cell lines. For example, heritability for red blood cell distribution width is most strongly enriched in TF motifs from K562, an erythroid cell line, heritability for rheumatoid arthritis, an autoimmune condition, is strongly enriched within TF motifs in ChIP-seq peaks from GM12878, a B-cell line; and heritability for serum cholesterol level is most strongly enriched in TF motifs from HepG2, a hepatocyte cell line (**Figure 5D**). These models are available for download through Factorbook for application to the summary statistics of additional genome-wide association studies (GWAS); we provide a Docker image and associated scripts for running this analysis through GitHub.

### High-resolution nucleosome and epigenetic profiles around binding sites

In the previous iteration of Factorbook, we generated aggregated epigenetic signal profiles, including histone modifications and nucleosome positions from MNase-seq, around the summits of TSS-proximal and TSS-distal transcription factor ChIP-seq peaks. We find that aggregating around motif instances rather than peak summits improves the resolution and phasing of epigenetic signals; additionally, it offers a natural orientation which reveals asymmetries in the organization of features around regulatory sites which have previously been suggested to be of biological relevance (26); we highlight, for example, asymmetric positioning of nucleosomes assayed by MNase-seq around oriented CTCF motif sites (**Figure 6A**). Therefore, on each factor’s page, Factorbook now displays aggregated signal profiles around motif instances for all cataloged motifs in addition to profiles surrounding ChIP-seq peak summits (illustrated for histone marks H3K4me3 and H3K4me1 around GATA4 motif sites, **Figure 6B**); we separate motif sites according to TSS proximity, which highlights differences in epigenetic profiles around TSS-proximal and TSS-distal sites.

**Figure 6.**
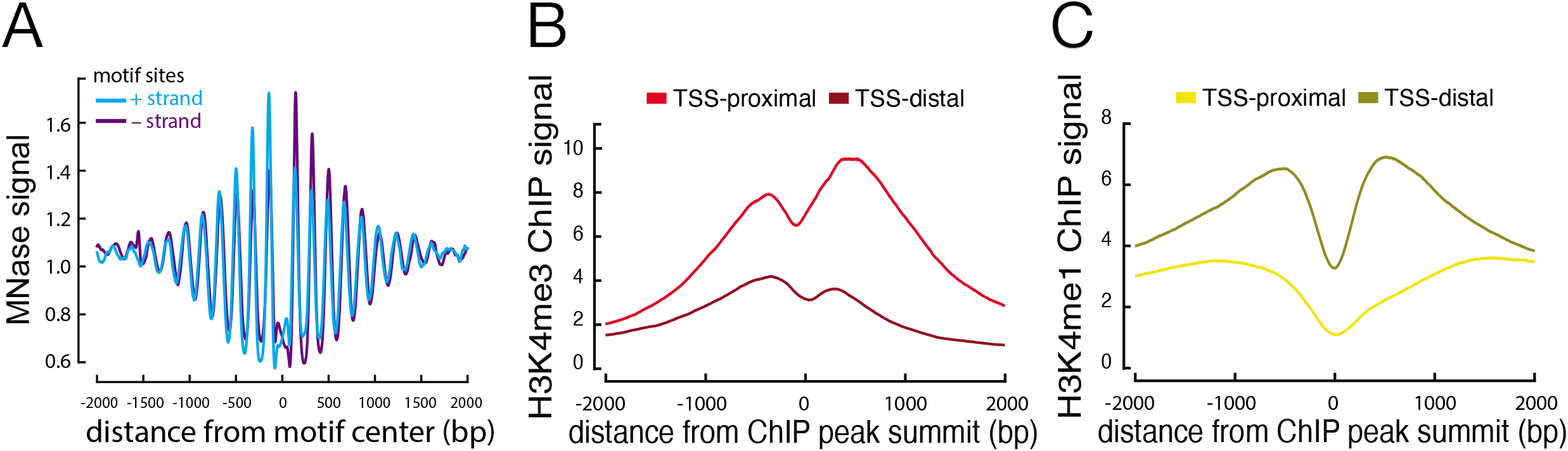
Epigenetic signal aggregation profiles on Factorbook. (**A**) Aggregated MNase-seq signal around CTCF motifs, highlighting asymmetry depending on motif orientation. (**B**) Aggregated H3K4me3 ChIP-seq signal around GATA4 peak summits as displayed in Factorbook. (**C**) Aggregated H3K4me1 ChIP-seq signal around GATA4 peak summits as displayed in Factorbook.

### Tools for machine learning and integrative analysis

Building deep learning models which can predict regulatory readouts is a primary focus of ongoing computational efforts in regulatory genomics. Prediction targets include cross-cell type transcription factor binding (13, 19) as well as epigenetic sequence profiles in a given cell type (12). Frequently these models include one-dimensional convolutional neural network layers which learn predictive sequences including transcription factor motifs. Transfer learning, or using existing models as starting points for new models applied to new tasks, has been proposed for sequence-based problems in biology (27, 28); seeding new models with our motif features could offer great potential to reduce training time while improving both the predictive power and human interpretability of learned models. Therefore we aimed to make Factorbook a platform for sharing motif kernels which can be applied directly to new machine learning tasks. To aid users in applying the kernels learned by our neural networks from HT-SELEX and ChIP-seq data to new biological questions, we provide the option to export all ZMotif-derived motifs in HDF5 format. These kernels may then be loaded into Python and used to seed weights in convolutional layers in a variety of commonly-used machine learning packages including PyTorch and Tensorflow. For users interested in more conventional downstream analysis, we also offer the option to export all MEME- and ZMotif-derived motifs as PWMs in MEME format, which may then be used by a variety of downstream tools including those in the MEME suite (20).

### Genomic visualization of motifs and TF binding sites

Human interaction remains essential in interpreting the biological significance of transcription factor motifs and regulatory elements. We implemented lightweight embedded genome visualizations within Factorbook which display TF peaks from ENCODE datasets alongside motif instances from our resource. Evolutionary conservation and relevant epigenetic signal profiles are displayed alongside gene and transcript tracks (**Figure 7A**). Additionally, we have designed a novel sequence importance track which scales bases in the reference sequence according to a signal track of associated scores; we demonstrate the use of this track to highlight evolutionarily conserved motif instances using PhyloP as the scaling score (**Figure 7B**). We have engineered this track to extend easily to additional scores provided through BigWig format signal tracks. All these tracks are rendered with vector graphics, allowing rich interactive interactions; for example, we layer popup views of SNPs and consensus logos onto these motif tracks in response to mouse events to aid users in quantifying the impact of alternative alleles using our LogoJS package. In addition, we have designed a public Factorbook trackhub for release on the UCSC Genome Browser (29).

**Figure 7.**
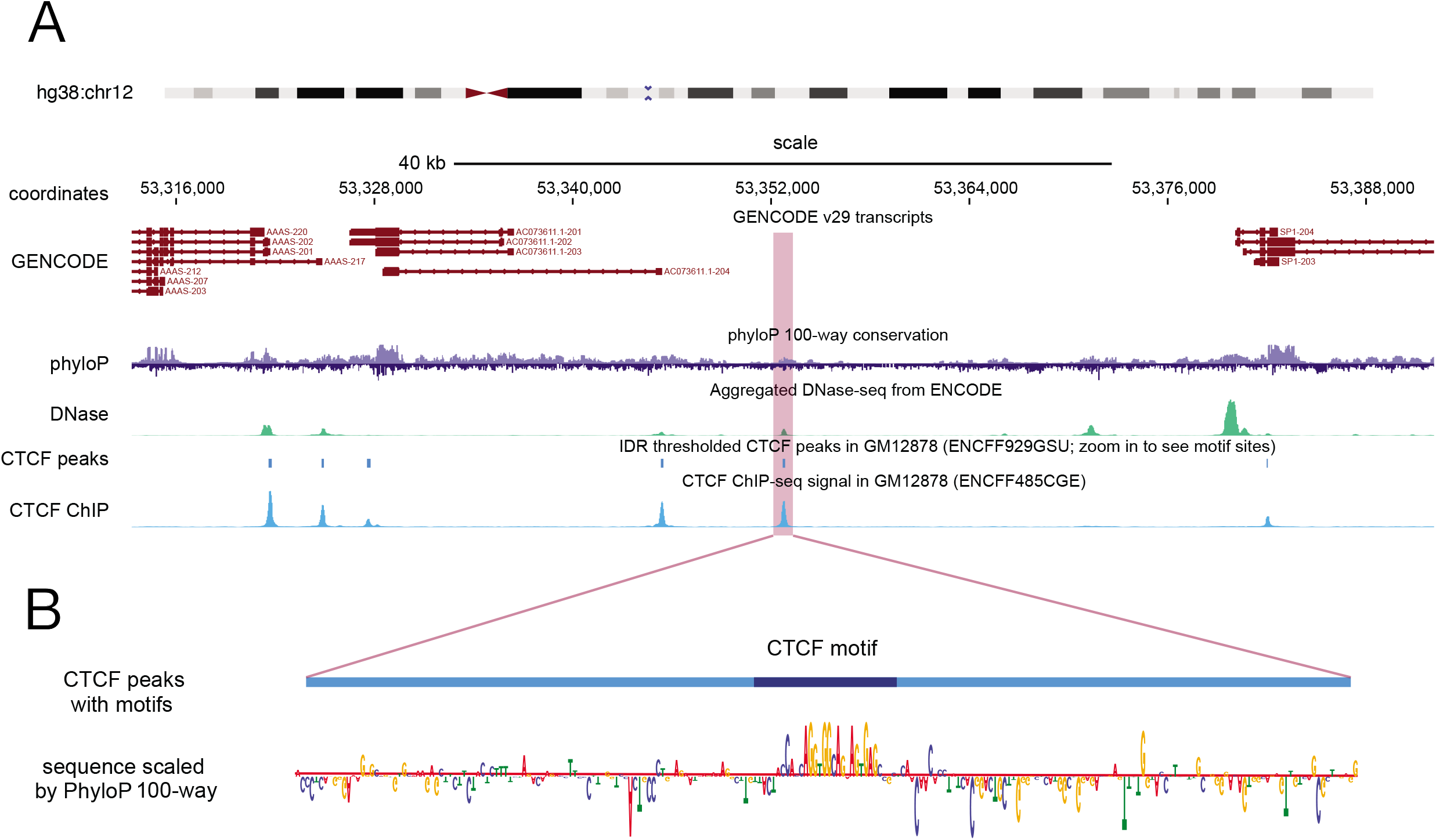
The Factorbook embedded genome browser view. (**A**) For a given experiment, ChIP-seq signal and IDR peaks from the ENCODE portal are displayed alongside transcripts and evolutionary conservation. (**B**) When the view is zoomed in, motif sites from the Factorbook catalog are displayed along with the underlying sequence scaled according to evolutionary conservation using a novel sequence importance track.

### Planned future expansion

We will further expand Factorbook to include integrative analysis of ENCODE Phase IV ChIP-seq, DNase-seq, and ATAC-seq data as they are released, and will update our motif and rDHS catalog accordingly. Additionally, we will update our motif instance catalog with the release of the final version of the ENCODE Registry of cCREs at the conclusion of ENCODE Phase IV.

## METHODS

### Motif discovery with MEME

We downloaded replicated IDR peaks from the ENCODE Portal for all non-histone ChIP-seq experiments in human and mouse from ENCODE Phases II and III. A full list of experiments and corresponding peak files is provided in **Supplement 1**. We then resized all peaks to a uniform 300 bp centered on the peak summit as provided in the peak file, sorted peaks by q-value, took the top 500, converted coordinates to FASTA sequence using the GRCh38 reference genome sequence and twoBitToFa tool from UCSC (1), and performed MEME (2) with the following parameters: *-zoops -dna -mod -nmotifs 5 -minw 6 -maxw 30 -revcomp*. The pipeline is available with a corresponding Docker image on GitHub (https://www.github.com/weng-lab/motif-workflow).

### Comparison with existing databases

We downloaded the full set of HOCOMOCO (3) and JASPAR 2020 (4, 5) motifs in MEME format. We then identified best matches for each of our HT-SELEX motifs and each of our MEME motifs using TOMTOM (2) with default parameters. The pipeline is available with a corresponding Docker image on GitHub (https://www.github.com/weng-lab/motif-workflow).

### Motif site identification within ChIP-seq peaks

We converted the original peak coordinates for each IDR peak set to a FASTA using the GRCh38 reference genome sequence and twoBitToFa tool from UCSC (1). We then scanned these sequences for each MEME motif using FIMO (2) with default parameters. For each motif instance, we then computed a single peak centrality value as the distance between the peak summit and the motif site center. Negative values indicate a motif upstream of the peak summit (toward a smaller genomic coordinate) for plus-strand motifs and downstream (toward a larger coordinate) for minus strand motifs. The pipeline is available with a corresponding Docker image on GitHub (https://www.github.com/weng-lab/motif-workflow).

### Conservation and DNase-seq aggregation

For each motif, we computed conservation and DNase-seq aggregate plots for all ChIP-seq peak motif sites. For conservation, we used the pyBigWig package to calculate the phyloP 100-way conservation at base-pair resolution for 1,000 base pairs in each direction centered on the motif site center; aggregation was strand-aware (directionality was reversed for minus strand motifs). For DNase-seq, we used the pysam package to count 5’ cut sites at base pair resolution within a 1,000 base pair window in each direction centered on the motif site center, also reversing orientation for minus-strand motifs. The pipeline is available with a corresponding Docker image on GitHub (https://www.github.com/weng-lab/motif-workflow).

### Motif site identification within rDHSs

We obtained representative DNase hypersensitive sites (rDHSs) from V3 of the ENCODE Registry of candidate Cis-Regulatory Elements (cCREs) (6) from the ENCODE Portal (accession ENCSR890YQQ) and converted the rDHSs to a FASTA using the GRCh38 reference sequence and twoBitToFa. We then scanned these sequences for each MEME motif passing QC using FIMO (2) with default parameters except for --max-stored-scores, which we set to 10 million given the larger size of the rDHS set. We then filtered the resulting motif sites at three different FIMO p-value thresholds: 10^−6^, 10^−7^, and 10^−8^.

### Motif UMAP

#### Distance metric

For each motif pair, we compute a Euclidean distance between position weight matrices for all possible alignments between the motifs, once with both oriented forward and once with one motif reverse complemented. For each aligned position, we compute the sum of the squared differences in information content for **A** in motif 1 vs. **A** in motif 2, **C** in motif 1 vs. **C** in motif 2, etc. When one motif extends past the end of the other, we consider the second motif to have 0 information content for each of **A**, **C**, **G**, and **T** at any extra positions. We take the square root of the summed differences at each position and normalize by dividing the total by the length of the shorter motif. The final distance between motif 1 and motif 2 is the minimum distance across all alignments, i.e., the distance at the optimal alignment. This distance metric is used both for UMAP and motif searching on Factorbook.

#### UMAP

We precomputed a distance matrix for all MEME motifs passing QC and a second distance matrix for all HT-SELEX motifs (7) using our Euclidean distance metric described above. We then performed UMAP on these distance matrices using the umap-learn package in Python with the following hyperparameters: min_dist=0.1, n_neighbors=5, 6, 7, 8, 9, or 10. For plotting, we colored motifs according to the DNA binding domain family for the corresponding factor as annotated by Lambert and colleagues (8).

### ZMotif: A convolution neural network for identifying transcription factor binding sites in TF ChIP-seq peaks

#### Network architecture

An input DNA sequence is one-hot-encoded (A=[1,0,0,0], C=[0,1,0,0], G=[0,0,1,0] and T=[0,0,0,1]). Sequences are padded with stretches of Ns ([0.25, 0.25. 0.25, 0.25]) on either side, equal in length to the width of the convolution kernels. If sequences are of variable length, they are padded with Ns to the length of the longest sequence. The reverse complement of the encoded sequence is generated and passed along with the original sequence to a shared convolution layer consisting of 16 kernels of width 24 and a linear activation function. Training a single, shared convolution layer on both the forward and reverse complement representations prevents the learning of averaged and duplicated motif representations. The bias of each convolution kernel is set to 0 so only the product of the convolution kernel and the one-hot-encoded sequence is passed to subsequent layers. The convolved sequence is then passed to two max pooling layers; the first over the strand dimension and the second over the spatial dimension. The maximum value over both dimensions for each kernel is then passed to a single output neuron with a sigmoid activation. The resulting logistic regression architecture, lacking intermediate layers, prevents the learning of distributed motif representations across multiple kernels. The weights of the output neuron are constrained to be greater than 0 to prevent the network from learning anti-motifs (i.e., motifs enriched in the negative sequence set compared to the positive set). The architecture is implemented and trained in Keras with the Model API and Tensorflow as the backend.

#### Convolution kernel k-mer initialization

A single convolution kernel is initialized with the one-hot-encoded representation of the most enriched 6-mer in input sequences. Initializing a single convolution kernel using the most strongly enriched k-mer often ensures that the corresponding motif is learned in full, especially for TFs with longer recognition sequences. All 6-mers in the input sequence are counted. The most significant k-mer is determined using a two-proportion z-test under the null hypothesis that the proportions of each individual k-mer are equal. The most enriched 6-mer is one-hot-encoded and inserted into the center of the first convolution kernel with the remaining values generated at random according to a uniform distribution.

#### Network training

Input sequences are randomly shuffled with 10% held out to monitor model accuracy and assess motif significance post-training. Sequences are fed to the network with a batch size of 32. If the classes are imbalanced, an equal number of positive and negative sequences (generated by dinucleotide shuffling positive sequences) are drawn in each batch. The model is trained using the Adam optimizer and cyclical learning rates between 0.1 and 0.01. The simple network architecture and small batch size permit the use of relatively high learning rates without experiencing exploding gradients. The network is trained for 1000 epochs consisting of 5000 total sequences. Fixing the number of epochs and sequences per epoch allows for a consistent training time independent of input datasets size. Stochastic weight averaging is used to create an ensemble model of the last 200 epochs.

#### Motif discovery from convolution kernels

Input positive sequences (HT-SELEX reads) are scanned simultaneously with all trained convolution kernels. If a given subsequence produces an activation greater than 0 (i.e., under our network constraints must contribute positively to the probability that the given sequence is bound by the TF), it is considered an instance of the corresponding kernel (also called a TF binding site). All instances of a given kernel are stacked and summed to produce a position frequency matrix and the corresponding position probability matrix (PPM) and position weight matrix (PWM). Motifs are trimmed by removing all positions from each end having an information content less than 0.25.

#### Assessing motif significance

For each convolution kernel, all output weights corresponding to every other kernel are set to 0. The area under the receiver-operator curve (auROC) of the resulting single feature logistic regression evaluated on the held out sequence set is used as a measure of the discovered motif’s significance.

### LD Score Regression

We downloaded summary statistics for GWAS for three distinct traits from the UK Biobank: red blood cell distribution width, rheumatoid arthritis, and serum cholesterol level. We used bedtools merge to generate union sets of ChIP-seq peaks for seven ENCODE cell lines: K562, GM12878, HepG2, MCF-7, H1-hESC, A549, and HEK293, by combining all IDR peaks from each of these cell lines. We then used bedtools intersect and bedtools subtract to identify motif sites and non-motif site sequences, respectively, within these peaks. We lifted both sets of regions from GRCh38 down to hg19 for compatibility with the summary statistics. We also lifted down the complete set of ChIP-seq motif sites as well as each set of union ChIP-seq peaks without motif sites removed.

We built a custom Docker image with all dependencies required for LDSC pre-installed (available for use through GitHub; see https://www.github.com/weng-lab/ldr). We then built LD score regression models for each set of lifted down regions by extending v2.2 of the LDSC baseline model (9) using the following command within the Docker image:

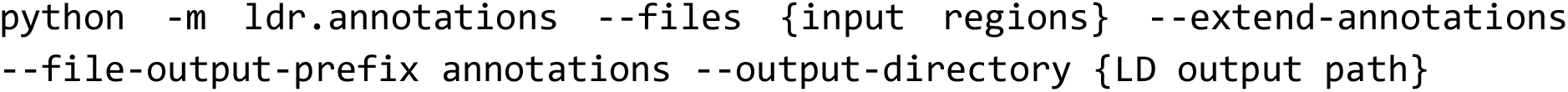

We packaged the outputs from this command into TAR archives for each region set. We then partitioned heritability for each set of summary statistics using the same Docker image and the following command:

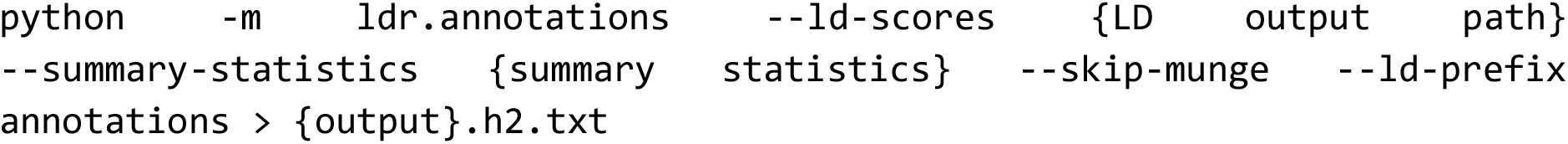

We then plotted heritability enrichment and error for each partition for each set of summary statistics.

## REFERENCES

1. Lambert,S.A., Jolma,A., Campitelli,L.F., Das,P.K., Yin,Y., Albu,M., Chen,X., Taipale,J., Hughes,T.R. and Weirauch,M.T. (2018) The Human Transcription Factors. Cell, 172, 650–665.

2. Jolma,A., Yan,J., Whitington,T., Toivonen,J., Nitta,K.R., Rastas,P., Morgunova,E., Enge,M., Taipale,M., Wei,G., et al. (2013) DNA-binding specificities of human transcription factors. Cell, 152, 327–339.

3. Johnson,D.S., Mortazavi,A., Myers,R.M. and Wold,B. (2007) Genome-wide mapping of in vivo protein-DNA interactions. Science, 316, 1497–1502.

4. Robertson,G., Hirst,M., Bainbridge,M., Bilenky,M., Zhao,Y., Zeng,T., Euskirchen,G., Bernier,B., Varhol,R., Delaney,A., et al. (2007) Genome-wide profiles of STAT1 DNA association using chromatin immunoprecipitation and massively parallel sequencing. Nat. Methods, 4, 651–657.

5. Kulakovskiy,I.V., Vorontsov,I.E., Yevshin,I.S., Sharipov,R.N., Fedorova,A.D., Rumynskiy,E.I., Medvedeva,Y.A., Magana-Mora,A., Bajic,V.B., Papatsenko,D.A., et al. (2018) HOCOMOCO: towards a complete collection of transcription factor binding models for human and mouse via large-scale ChIP-Seq analysis. Nucleic Acids Res., 46, D252–D259.

6. Fornes,O., Castro-Mondragon,J.A., Khan,A., van der Lee,R., Zhang,X., Richmond,P.A., Modi,B.P., Correard,S., Gheorghe,M., Baranašić,D., et al. (2020) JASPAR 2020: update of the open-access database of transcription factor binding profiles. Nucleic Acids Res., 48, D87–D92.

7. Newburger,D.E. and Bulyk,M.L. (2009) UniPROBE: an online database of protein binding microarray data on protein-DNA interactions. Nucleic Acids Res., 37, D77–82.

8. Weirauch,M.T., Yang,A., Albu,M., Cote,A.G., Montenegro-Montero,A., Drewe,P., Najafabadi,H.S., Lambert,S.A., Mann,I., Cook,K., et al. (2014) Determination and inference of eukaryotic transcription factor sequence specificity. Cell, 158, 1431–1443.

9. Wang,J., Zhuang,J., Iyer,S., Lin,X., Whitfield,T.W., Greven,M.C., Pierce,B.G., Dong,X., Kundaje,A., Cheng,Y., et al. (2012) Sequence features and chromatin structure around the genomic regions bound by 119 human transcription factors. Genome Res., 22, 1798–1812.

10. Wang,J., Zhuang,J., Iyer,S., Lin,X.-Y., Greven,M.C., Kim,B.-H., Moore,J., Pierce,B.G., Dong,X., Virgil,D., et al. (2013) Factorbook.org: a Wiki-based database for transcription factor-binding data generated by the ENCODE consortium. Nucleic Acids Res., 41, D171–6.

11. Quang,D. and Xie,X. (2019) FactorNet: A deep learning framework for predicting cell type specific transcription factor binding from nucleotide-resolution sequential data. Methods, 166, 40–47.

12. Avsec,Ž., Weilert,M., Shrikumar,A., Krueger,S., Alexandari,A., Dalal,K., Fropf,R., McAnany,C., Gagneur,J., Kundaje,A., et al. (2021) Base-resolution models of transcription-factor binding reveal soft motif syntax. Nat. Genet., 53, 354–366.

13. Alipanahi,B., Delong,A., Weirauch,M.T. and Frey,B.J. (2015) Predicting the sequence specificities of DNA- and RNA-binding proteins by deep learning. Nat. Biotechnol., 33, 831–838.

14. ENCODE Project Consortium, Moore,J.E., Purcaro,M.J., Pratt,H.E., Epstein,C.B., Shoresh,N., Adrian,J., Kawli,T., Davis,C.A., Dobin,A., et al. (2020) Expanded encyclopaedias of DNA elements in the human and mouse genomes. Nature, 583, 699–710.

15. Finucane,H.K., Bulik-Sullivan,B., Gusev,A., Trynka,G., Reshef,Y., Loh,P.-R., Anttila,V., Xu,H., Zang,C., Farh,K., et al. (2015) Partitioning heritability by functional annotation using genome-wide association summary statistics. Nat. Genet., 47, 1228–1235.

16. Khan,A., Fornes,O., Stigliani,A., Gheorghe,M., Castro-Mondragon,J.A., van der Lee,R., Bessy,A., Chèneby,J., Kulkarni,S.R., Tan,G., et al. (2018) JASPAR 2018: update of the open-access database of transcription factor binding profiles and its web framework. Nucleic Acids Res., 46, D260–D266.

17. Mathelier,A., Fornes,O., Arenillas,D.J., Chen,C.-Y., Denay,G., Lee,J., Shi,W., Shyr,C., Tan,G., Worsley-Hunt,R., et al. (2016) JASPAR 2016: a major expansion and update of the open-access database of transcription factor binding profiles. Nucleic Acids Res., 44, D110–5.

18. Hume,M.A., Barrera,L.A., Gisselbrecht,S.S. and Bulyk,M.L. (2015) UniPROBE, update 2015: new tools and content for the online database of protein-binding microarray data on protein-DNA interactions. Nucleic Acids Res., 43, D117–22.

19. Chen,C., Hou,J., Shi,X., Yang,H., Birchler,J.A. and Cheng,J. (2021) DeepGRN: prediction of transcription factor binding site across cell-types using attention-based deep neural networks. BMC Bioinformatics, 22, 38.

20. Bailey,T.L., Johnson,J., Grant,C.E. and Noble,W.S. (2015) The MEME Suite. Nucleic Acids Res., 43, W39–49.

21. Becht,E., McInnes,L., Healy,J., Dutertre,C.-A., Kwok,I.W.H., Ng,L.G., Ginhoux,F. and Newell,E.W. (2018) Dimensionality reduction for visualizing single-cell data using UMAP. Nat. Biotechnol., 10.1038/nbt.4314.

22. McInnes,L., Healy,J. and Melville,J. (2018) UMAP: Uniform Manifold Approximation and Projection for Dimension Reduction. arXiv:1802.03426 [cs, stat].

23. Gupta,S., Stamatoyannopoulos,J.A., Bailey,T.L. and Noble,W.S. (2007) Quantifying similarity between motifs. Genome Biol., 8, R24.

24. Yin,Y., Morgunova,E., Jolma,A., Kaasinen,E., Sahu,B., Khund-Sayeed,S., Das,P.K., Kivioja,T., Dave,K., Zhong,F., et al. (2017) Impact of cytosine methylation on DNA binding specificities of human transcription factors. Science, 356.

25. Pollard,K.S., Hubisz,M.J., Rosenbloom,K.R. and Siepel,A. (2010) Detection of nonneutral substitution rates on mammalian phylogenies. Genome Res., 20, 110–121.

26. Kundaje,A., Kyriazopoulou-Panagiotopoulou,S., Libbrecht,M., Smith,C.L., Raha,D., Winters,E.E., Johnson,S.M., Snyder,M., Batzoglou,S. and Sidow,A. (2012) Ubiquitous heterogeneity and asymmetry of the chromatin environment at regulatory elements. Genome Res., 22, 1735–1747.

27. Mignone,P., Pio,G., D’Elia,D. and Ceci,M. (2020) Exploiting transfer learning for the reconstruction of the human gene regulatory network. Bioinformatics, 36, 1553–1561.

28. Heinzinger,M., Elnaggar,A., Wang,Y., Dallago,C., Nechaev,D., Matthes,F. and Rost,B. (2019) Modeling aspects of the language of life through transfer-learning protein sequences. BMC Bioinformatics, 20, 723.

29. Kent,W.J., Sugnet,C.W., Furey,T.S., Roskin,K.M., Pringle,T.H., Zahler,A.M. and Haussler,D. (2002) The human genome browser at UCSC. Genome Res., 12, 996–1006.

## References

0. Kent,W.J., Sugnet,C.W., Furey,T.S., Roskin,K.M., Pringle,T.H., Zahler,A.M. and Haussler,D. (2002) The human genome browser at UCSC. Genome Res., 12, 996–1006.

1. Bailey,T.L., Boden,M., Buske,F.A., Frith,M., Grant,C.E., Clementi,L., Ren,J., Li,W.W. and Noble,W.S. (2009) MEME SUITE: tools for motif discovery and searching. Nucleic Acids Res., 37, W202–8.

2. Kulakovskiy,I.V., Vorontsov,I.E., Yevshin,I.S., Sharipov,R.N., Fedorova,A.D., Rumynskiy,E.I., Medvedeva,Y.A., Magana-Mora,A., Bajic,V.B., Papatsenko,D.A., et al. (2018) HOCOMOCO: towards a complete collection of transcription factor binding models for human and mouse via large-scale ChIP-Seq analysis. Nucleic Acids Res., 46, D252–D259.

3. Khan,A., Fornes,O., Stigliani,A., Gheorghe,M., Castro-Mondragon,J.A., van der Lee,R., Bessy,A., Chèneby,J., Kulkarni,S.R., Tan,G., et al. (2018) JASPAR 2018: update of the open-access database of transcription factor binding profiles and its web framework. Nucleic Acids Res., 46, D260–D266.

4. Fornes,O., Castro-Mondragon,J.A., Khan,A., van der Lee,R., Zhang,X., Richmond,P.A., Modi,B.P., Correard,S., Gheorghe,M., Baranašić,D., et al. (2020) JASPAR 2020: update of the open-access database of transcription factor binding profiles. Nucleic Acids Res., 48, D87–D92.

5. ENCODE Project Consortium, Moore,J.E., Purcaro,M.J., Pratt,H.E., Epstein,C.B., Shoresh,N., Adrian,J., Kawli,T., Davis,C.A., Dobin,A., et al. (2020) Expanded encyclopaedias of DNA elements in the human and mouse genomes. Nature, 583, 699–710.

6. Jolma,A., Yan,J., Whitington,T., Toivonen,J., Nitta,K.R., Rastas,P., Morgunova,E., Enge,M., Taipale,M., Wei,G., et al. (2013) DNA-binding specificities of human transcription factors. Cell, 152, 327–339.

7. Lambert,S.A., Jolma,A., Campitelli,L.F., Das,P.K., Yin,Y., Albu,M., Chen,X., Taipale,J., Hughes,T.R. and Weirauch,M.T. (2018) The Human Transcription Factors. Cell, 172, 650–665.

8. Finucane,H.K., Bulik-Sullivan,B., Gusev,A., Trynka,G., Reshef,Y., Loh,P.-R., Anttila,V., Xu,H., Zang,C., Farh,K., et al. (2015) Partitioning heritability by functional annotation using genome-wide association summary statistics. Nat. Genet., 47, 1228–1235.

